# Elucidating the functional role of predicted miRNAs in post-transcriptional gene regulation along with symbiosis in *Medicago truncatula*

**DOI:** 10.1101/346858

**Authors:** Roy Chowdhury Moumita, Jolly Basak, Ranjit Prasad Bahadur

## Abstract

Non-coding RNAs (ncRNAs) are found to be important regulator of gene expression because of their ability to modulate post-transcriptional processes. microRNAs are small ncRNAs which inhibit translational and post-transcriptional processes whereas long ncRNAs are found to regulate both transcriptional and post-transcriptional gene expression. *Medicago truncatula* is a well-known model plant for studying legume biology and is also used as a forage crop. In spite of its importance in nitrogen fixation and soil fertility improvement, little information is available about Medicago ncRNAs that play important role in symbiosis. To understand the role of Medicago ncRNAs in symbiosis and regulation of transcription factors, we have identified novel miRNAs and tried to establish an interaction model with their targets. 149 novel miRNAs are predicted along with their 770 target proteins. We have shown that 51 of these novel miRNAs are targeting 282 lncRNAs. We have analyzed the interactions between miRNAs and their target mRNAs as well as their targets on lncRNAs. Role of Medicago miRNAs in the regulation of various transcription factors were also elucidated. Knowledge gained from this study will have a positive impact on the nitrogen fixing ability of this important model plant, which in turn will improve the soil fertility.

## INTRODUCTION

Functional RNA molecules, encoded by non-coding RNA genes, play important roles in regulating the expression of various genes (1). MicroRNAs (miRNAs), first discovered in *Caenorhabditis elegans* (lin-4) (2) are 20-24 nucleotides long, endogenous, small non-coding ribonucleic acids that regulate gene expression post-transcriptionally in plants, animals and viruses (3,4). miRNAs regulate gene expression in various ways including cleavage, degradation and decaying of mRNA and nascent polypeptide, inhibition of translation by forming miRNA-mRNA complex, ribosome drop-off resulting in premature termination, sequestration of target mRNAs in P-bodies that are distinct foci consisting enzymes involved in mRNA turnover (5), transcriptional inhibition by miRNA mediated chromatin reorganization, inhibition of translation initiation by suppressing the cap recognition by elF4E and 60S-40S joining (6–16). Plant miRNAs can repress translation by binding in a near-perfect manner with target mRNAs (17) where animal miRNAs can recognize target mRNAs through six to eight nucleotides long seed region of the 5’end of the mRNA (18).

Various miRNAs have been discovered in animals, plants and viruses with the help of conventional experimental strategies along with bioinformatics tools (19,20). In spite of present state-of-the art technologies including high throughput sequencing, identification of miRNAs in various organisms is still complex and need proper blending of experimental strategies and computational approaches to get more accurate results. miRNAs can be identified by computational approaches based on the presence of hairpin loop in pre-miRNA secondary structures, although stem-loop structure is not an unique feature of pre-miRNA as it is present in other RNAs as well (21–23). To resolve this problem, minimum folding free energy index (MFEI) and AU content were also considered along with secondary structures to distinguish pre-miRNAs from the other RNAs. MFEI and AU content of miRNAs are significantly higher than other ncRNAs (24). Normalized base-pairing propensity, normalized Shannon entropy, normalized base-pair distance and degree of compactness (25,26) were also included in recent computational based identification of miRNAs along with normalized MFEI. In addition simple sequence repeats (SSRs), being an important component of pre-miRNAs, was successfully applied to remove the false positives in the prediction (26–30).

*Medicago truncatula*, commonly known as barrelclover is a small legume found in the Mediterranean region (31). *M. truncatula* is routinely used in genomic research as a model organism to study the legume biology owing to its small diploid genome, self-fertility, rapid generation time and formation of symbiosis with rhizobia (32). *M. truncatula* genome contains nine symbiotic leghaemoglobins, which is more than twice the number of leghaemoglobins present in *Glycine max* and *Lotus japonicas* (32). Presence of many nodulation related genes and the leghaemoglobins in Medicago genome (32) made this plant useful for nitrogen fixation to improve soil fertility. *M. truncatula,* being an important forage crop, is harvested as winter annual crop in the Mediterranean regions, and can also be harvested as short seasonal annual crops in fall and summer in north-central United States of America (USA) (33). It has also shown potentiality as an emergency forage crop in northern USA (34). In spite of the presence of large number of Medicago miRNAs in miRBase21, limited studies have been done so far about the involvement of miRNAs in the regulation of symbiosis and transcription. Lack of this information prompted us to identify miRNAs that are linked with symbiosis and transcription. In this study we have predicted 149 novel miRNAs, which are not present in miRBase22 and establish a connection between these novel miRNAs with various transcription factors and proteins involved in symbiosis. We also have identified 770 mRNAs as their targets. In addition, we have analyzed the interactions between miRNAs and their mRNA targets as well as the interactions between miRNAs and their targets on lncRNAs. Owing to the important role of Medicago in symbiosis, a special emphasis was given to decipher the symbiosis related gene regulation through analyzing the interactions between predicted miRNAs and their nodulin target proteins. Along with that we have also elucidated diverse role of miRNAs in the regulation of various transcription factors of Medicago.

## Materials and Methods

### Data collection and preparation

We have downloaded 8495 mature miRNA sequences available from 73 species of Viridiplantae in miRBase21 (35) and used them for standard dataset preparation. *M. truncatula* genome sequences and coding DNA sequences were downloaded from Plant Genome and System Biology database (36). lncRNAs of *M. truncatula* were obtained from *Green Non-Coding Database* (37).

### Prediction of miRNAs in *M. truncatula*

For prediction of miRNAs, BLAST (38) search was performed by submitting the mature Viridiplantae miRNAs as query and the Medicago genome as the subject. All possible lengths of upstream and/or downstream sequences ranging between 55 and 505 nucleotides were extracted from the genome following Nithin, et al paper (26) and were aligned with the mature viridiplantae miRNAs. The sequences of BLAST output were retained with e-value cut off 1000, word size 7 and mismatch is less than 4 (39). In order to remove the protein coding sequences, an un-gapped BLASTX with the sequence identity cut-off ≥ 80 % was performed with all the extracted sequences as query and the protein sequences of *Medicago* as subject. After CDS removal, the obtained pre-miRNAs were subjected to the following criteria to predict mature miRNAs: (i) formation of stem-loop structure with MFEI ≥ 0.41, (ii) presence of a mature miRNA sequence in one arm of the stem-loop structure, (iii) maximum six mismatches between miRNA and miRNA* sequence, (iv) complete absence of any loop or break in miRNA* sequences. These four criteria must be satisfied by the predicted mature miRNAs. Along with these parameters, several other parameters namely AU content, normalized base pairing propensity (Npb), normalized base pairing distance (ND), normalized Shannon entropy (NQ) and SSRs were also taken into considerations and their cut-off values were set following Nithin et al., 2015 (26). The flowchart of the algorithm is shown in Fig 1.

**Fig 1:**
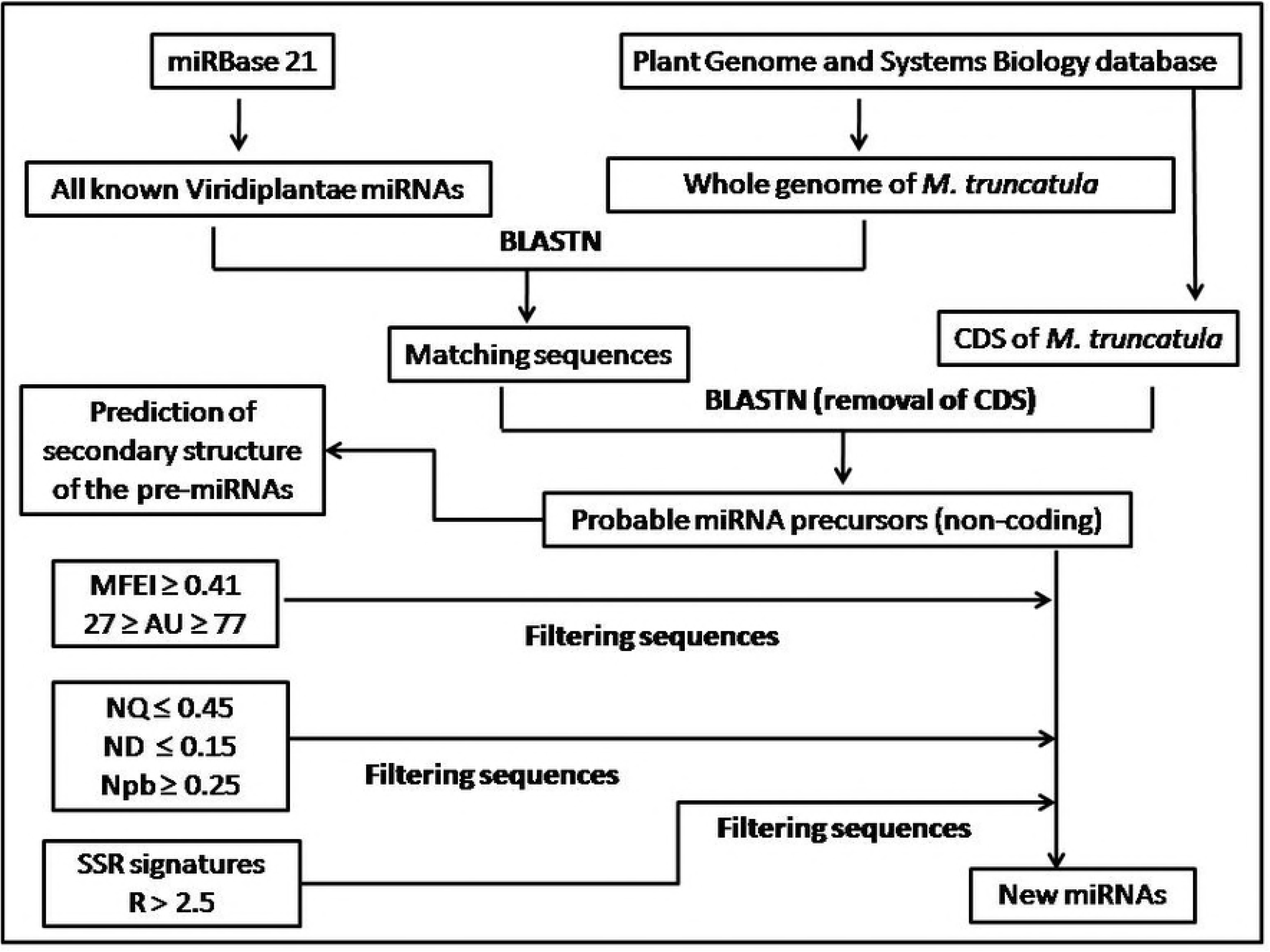
Flowchart for prediction of miRNAs in *M. truncatula*

Sequences satisfying all the above mentioned criteria were selected as mature miRNAs. Predicted miRNAs, which are absent in the miRBase21 are assigned as novel miRNAs.

### Target prediction and GO analysis of the predicted targets

Targets for the novel miRNAs were predicted using psRNATarget server by using *M. truncatula* transcript as subject and novel miRNAs as query. False predictions were removed by keeping 2.0 as a stringent value for expectation threshold according to Nithin et al. (2015) (26). The nucleotides for complementary scoring and hpsize were kept as same as the length of the mature miRNA. Maximum energy of unpairing the target site was set to 25 kcal. The flanking length around target site for target accessibility analysis was set between 17 bp upstream and 13 bp downstream (26). Sequence range of the central mismatch was adjusted with the varying miRNA length (26). Functional annotation of the target proteins were carried out using UniProt-GOA (40). Corresponding protein IDs of the target mRNAs and the UniProt *M. truncatula* proteome UP000002051 was used to obtain the information about the biological processes, molecular functions and the cellular components where the target proteins were found to be localized.

### Prediction of miRNA targets on lncRNAs

miRNA targets on lncRNAs were obtained using psRNATarget server by submitting the lncRNAs of *M. truncatula* as subject and mature miRNAs as query. The same conditions used to predict the mRNA targets were used here. Interaction network between the novel miRNAs and their target lncRNAs are shown using cytoscape.

## Results

### Prediction of new miRNAs in *M. truncatula*

BLAST search was performed using the whole genome of *M. truncatula* as subject and mature miRNAs as query. After removing the CDS, 746738 sequences are obtained, which are further examined for secondary structure, SSR, MFEI and the criteria described in materials and methods section (26). We obtained 297 sequences which satisfied all the criteria and are designated as putative pre-miRs. Finally from these putative pre-miRs, 178 mature non-redundant miRNAs are extracted, of which 149 miRNAs are absent in the miRBase21 and are designated as novel. List of 178 mature miRNAs along with the necessary criteria are provided in supplementary Table S1.

These 149 novel miRNAs belong to 40 different miRNA families (Table 1). Two families, miR5281 and miR5021, are found to be highly populated containing 41 and 21 members, respectively. Remaining families are less populated with number of members varying from one to nine. This distribution is in concordance with other species in Viridiplantae (41). Length of the predicted novel miRNAs varies from 16 to 24 nucleotides with an average length of 20 nucleotides (±2.7), while majority of the miRNAs have the length of 21 nucleotides. The distribution of the mature miRNA length is shown in Fig 2. The distribution of the miRNAs in the chromosome is shown in Fig 3.

**Fig 2:**
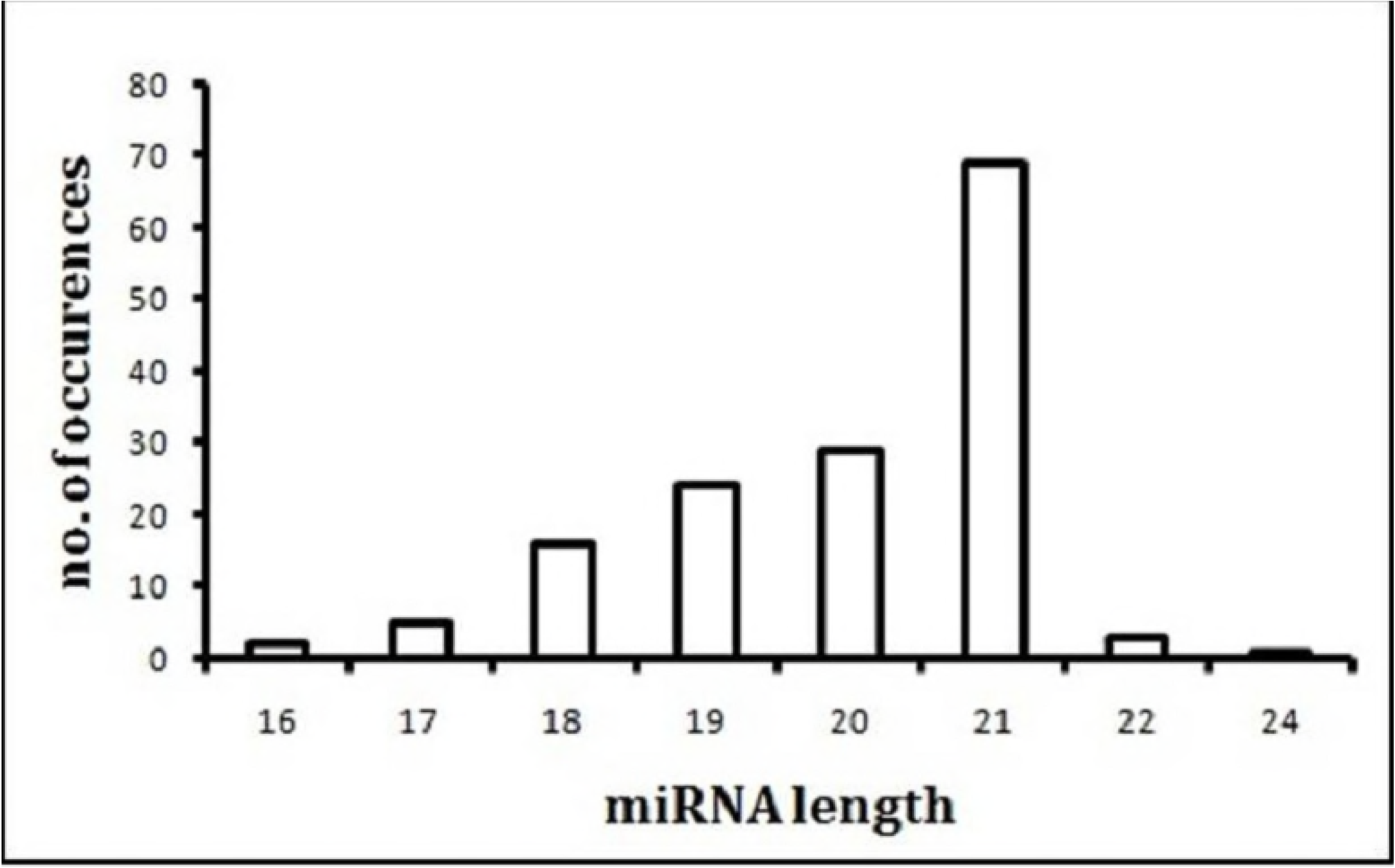
Distribution of length of predicted miRNAs of *M. truncatula L.*

**Fig 3:**
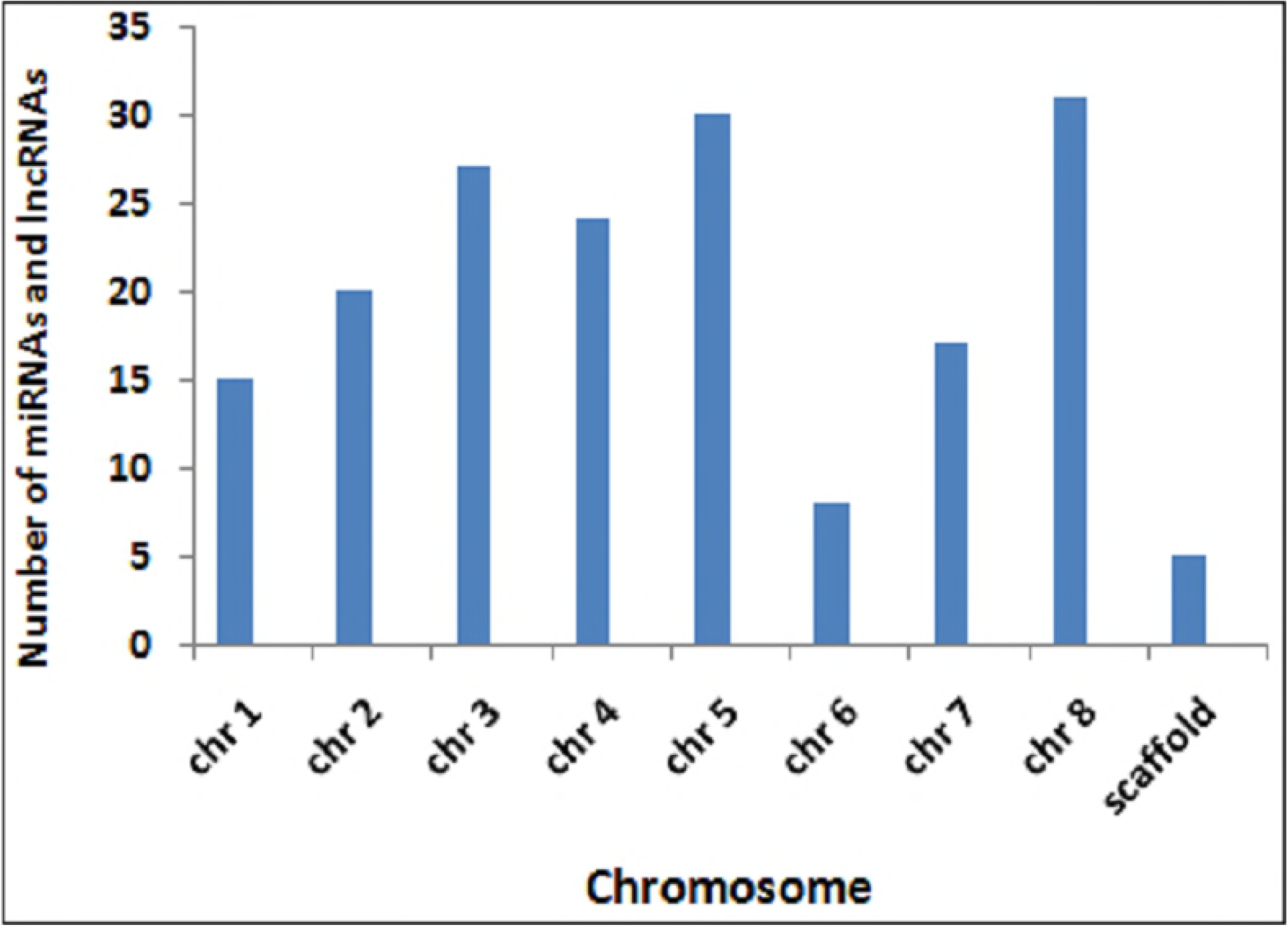
Chromosomal distribution of miRNAs and their target lncRNAs

**Table 1:** Distribution of miRNAs within different miRNA families of *M. truncatula L.*

| miRNA families | No. of members/family |
| --- | --- |
| miR5281 | 41 |
| miR5021 | 21 |
| miR414 | 9 |
| miR156 | 6 |
| miR172, miR845 | 5 |
| miR1533, miR169, miR5181 | 4 |
| miR171, miR5174, miR8155 | 3 |
| miR166, miR393, miR477, miR5650 | 2 |
| miR1171, miR1514, miR1530, miR160,<br>miR162, miR164, miR165, miR168, miR3445,<br>miR390, miR3933, miR398, miR407, miR4233,<br>miR4248, miR447, miR5014, miR5016,<br>miR5017, miR5171, miR5183, miR530,<br>miR5645, miR5649, miR5666, miR5998,<br>miR773, miR774, miR7745, miR829, miR837,<br>miR838, miR870 | 1 |

### Target prediction and GO analysis of the predicted targets

A total of 770 target proteins belong to 23 different families were obtained from psRNATarget server for 93 novel miRNAs. Among these targets, biological functions of 621 are known, while others are uncharacterized or hypothetical proteins. Interaction between miRNA and their targets are shown in Fig. 4a as a network model. Frequency distribution of these miRNAs and their targets are shown in Fig. 4b. Details of targets of the novel miRNAs are provided in supplementary table S2.

**Fig 4a:**
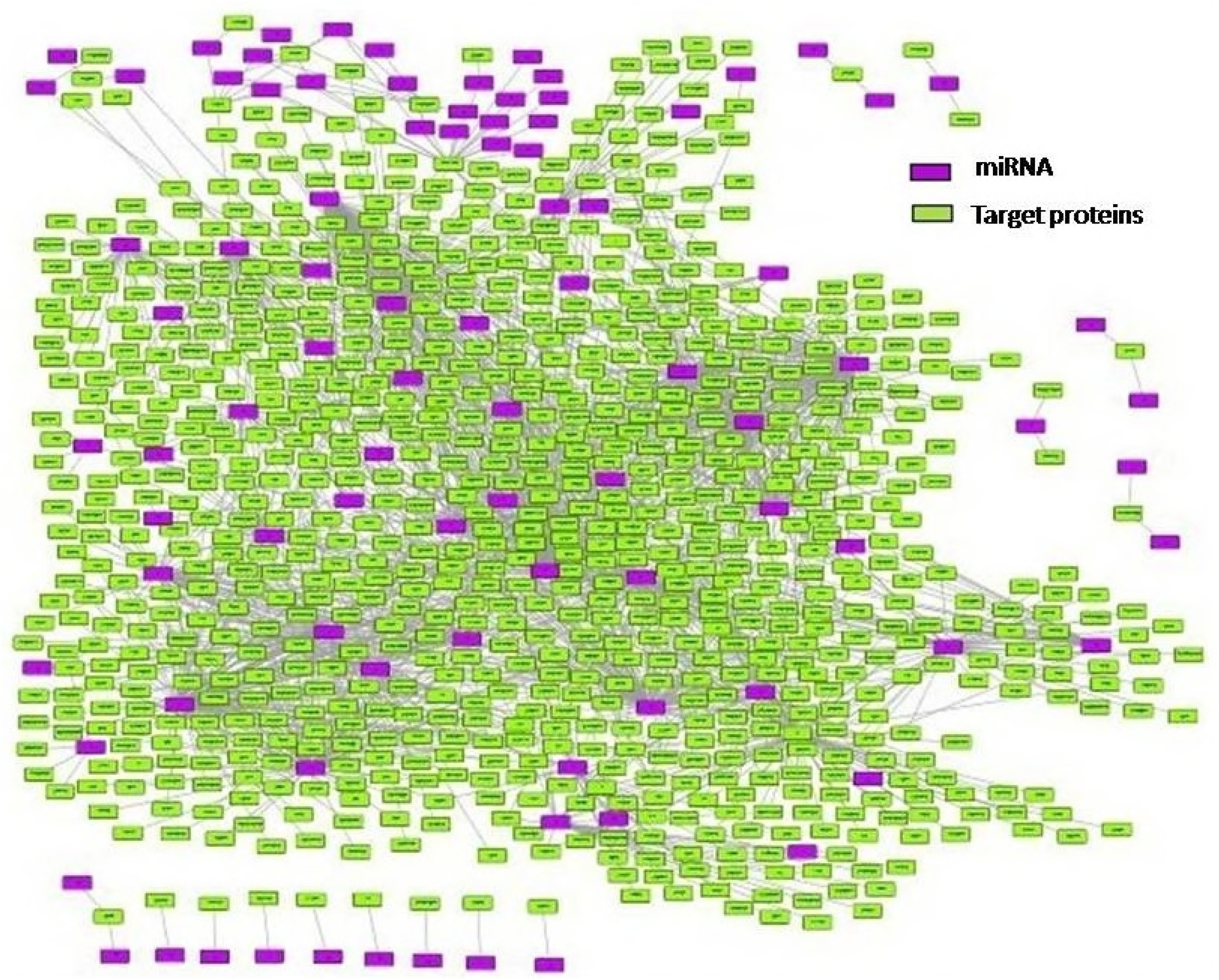
Network model of miRNAs and target proteins

**Fig 4b:**
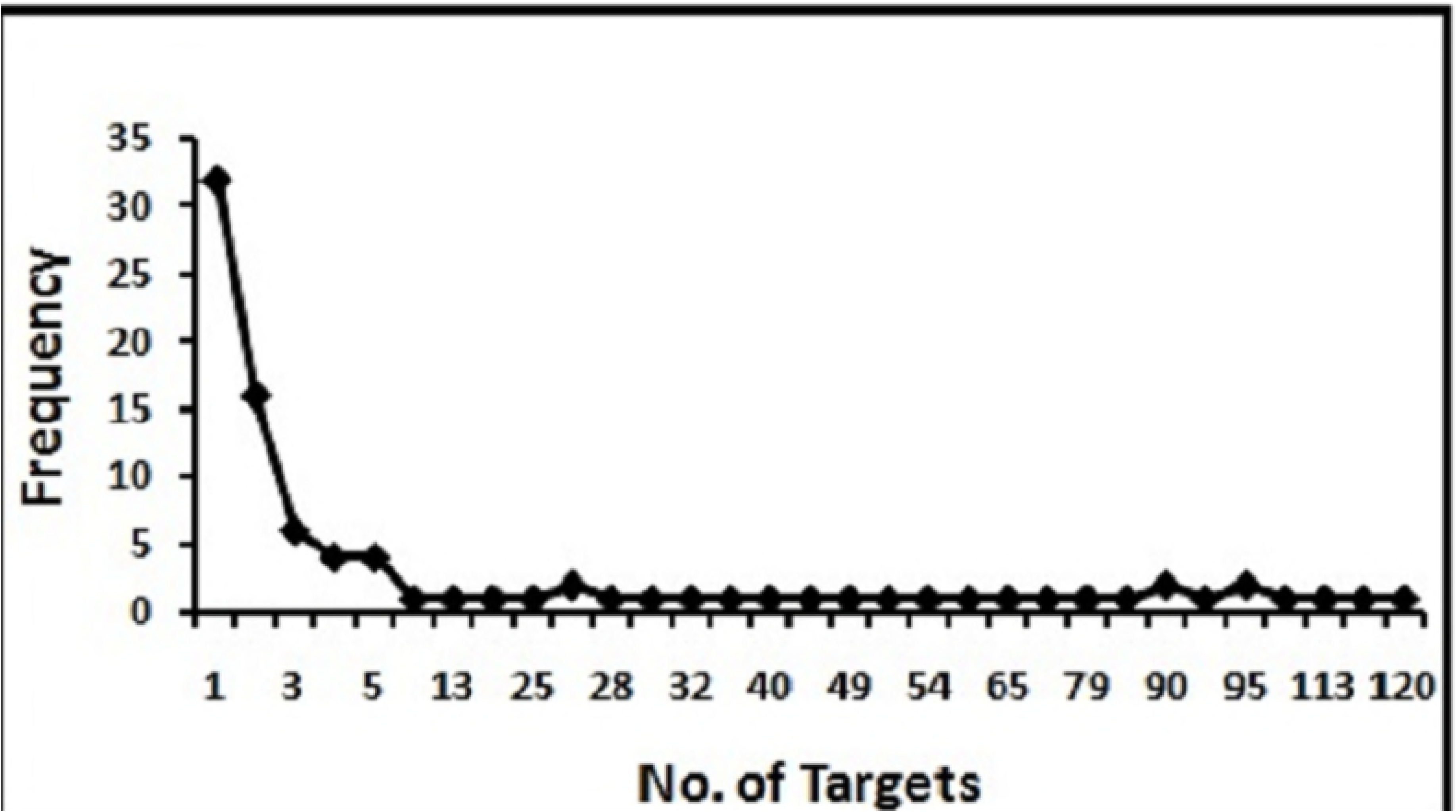
Frequency distribution of miRNA-target protein interaction

The number of targets of these miRNAs varies from one to 120. Among those, mtr-miR5021d (120 targets), mtr-miR5021c (118 targets) and mtr-miR5021s (113 targets) target large number of protein coding sequences. Majority of the targets code for various transporters, ribosomal proteins, receptors, kinases, helicases, transcription factors, trans-membrane proteins, nodulins and heme binding proteins. We find 32 miRNAs which target only one mRNA.

Among these predicted targets some are targeted by the same miRNA families of other Viridiplantae species. mtr-miR172a and mtr-miR172b both target AP2 domain transcription factor and AP2-like ethylene-responsive transcription factor, and is in accordance with the previous findings (41–43). Mtr-miR 166 targets class III homeodomain leucine zipper protein and bZIP transcription factor which is in agreement with the published reports (44–46).

Targets of the miRNAs also include various nodulins and heme binding proteins like legumin A2, non-symbiotic hemoglobin, nodulin MtN21/EamA-like transporter family protein, nodule inception protein, Nodule Cysteine-Rich (NCR) secreted peptide, nodulation receptor kinase-like protein and Nodule-specific Glycine Rich Peptide. miRNAs inhibit these target sequences either by translational repression or by cleavage. Multiple miRNAs target the same protein coding sequences, however the mode of inhibition being different. For an example, mtr-miR414c and mtr-miR5281h both target nodulin MtN21/EamA-like transporter family protein coding mRNA but mtr-miR414c inhibits by cleaving the sequence whereas mtr-miR5281h represses the translation of the protein coding sequence. *M. truncatula* is a well known model organism for the legume biology (32) and understanding the interactions between novel miRNAs and their nodulin targets will significantly enhance our knowledge on symbiosis. *M. truncatula* being a fabaceae plant, contains rhizobia within the nodules producing nitrogenous compounds that helps in growth and creating a forte of protection from other plants (47). During decay the fixed nitrogen get released and makes the nitrogen available for other plants, thereby improving the soil quality. Since the above mentioned miRNAs were found to target nodulin proteins, they in-turn are involved in the regulation of the nodule. For example, Nodule Cysteine-Rich (NCR) secreted peptide is important for nodule morphogenesis and nodulation receptor kinase-like protein is involved in the perception of symbiotic fungi and bacteria. A schematic representation of this symbiotic regulation is depicted in Fig 5. A network model is shown in this flowchart where the different miRNAs have been shown to target a specific protein involved in symbiosis.

**Fig 5:**
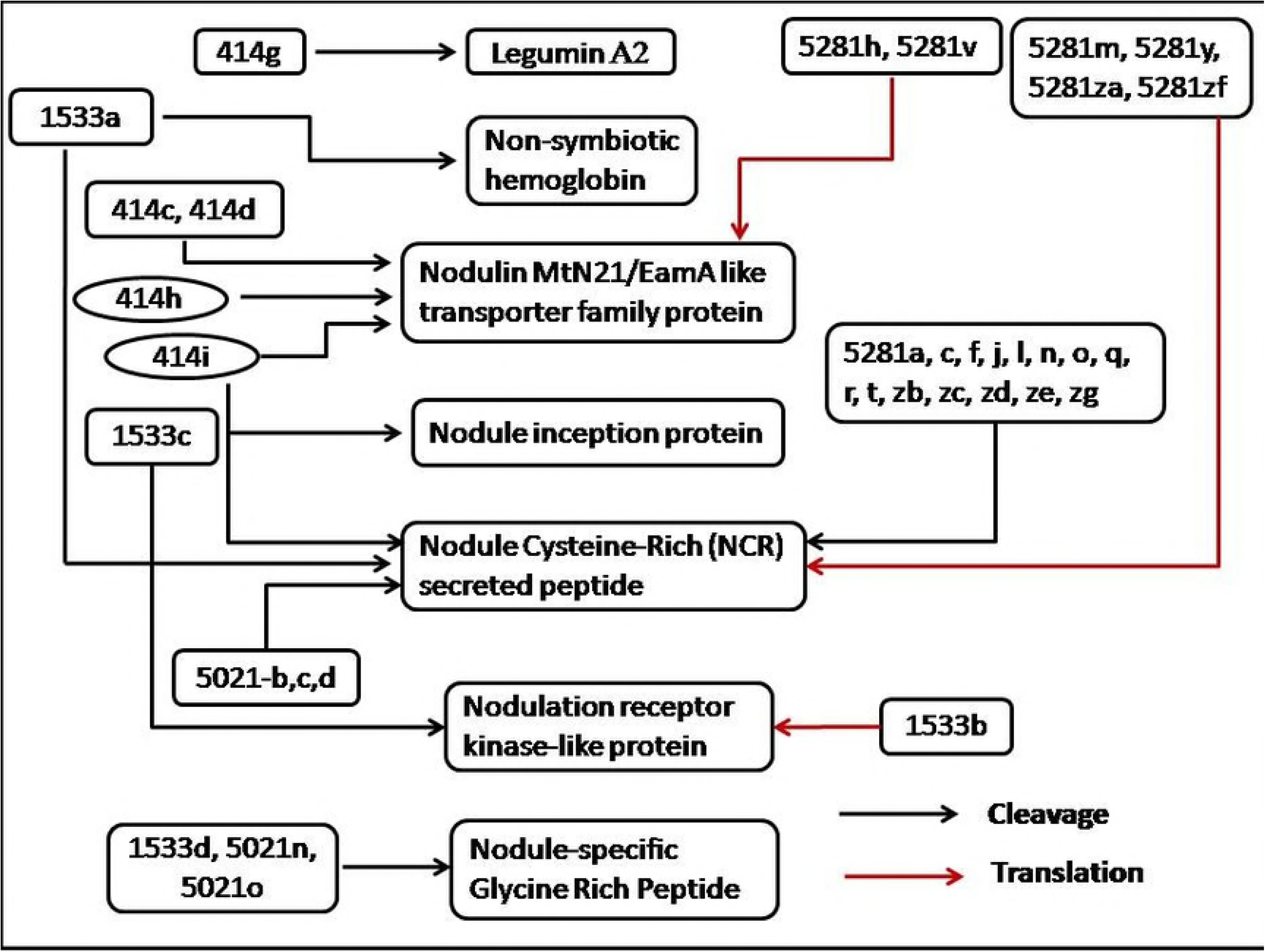
Symbiotic regulation of nodulin proteins by miRNAs

Many miRNAs target various transcription factors. mtr-miR166b, mtr-miR5021n, mtr-miR5021o and mtr-miR5021q target bZIP transcription factor. CCAAT-binding transcription factor was targeted by mtr-miR169b and mtr-miR169d, whereas, AP2 domain transcription factor was targeted by mtr-miR172a, mtr-miR172b, mtr-miR414c, mtr-miR414i and mtr-miR1533c. mtr-miR414a, mtr-miR414c and mtr-miR414i target heat shock transcription factor A3. myb DNA-binding domain protein and basic helix loop helix (bHLH) DNA-binding family protein are targeted by mtr-miR414b and mtr-miR414c, respectively. Different types of zinc finger proteins are targeted by mtr-miR414b, mtr-miR414c, mtr-miR414e, mtr-miR414g, mtr-miR414i, mtr-miR838, mtr-miR1533b, mtr-miR1533c, mtr-miR5014, mtr-miR5021b, mtr-miR5021c, mtr-miR5021d, mtr-miR5021g, mtr-miR5021h, mtr-miR5021i, mtr-miR5021l, mtr-miR5021m, mtr-miR5021n, mtr-miR5021o, mtr-miR5021q, mtr-miR5021s and mtr-miR5021u. MADS-box transcription factor is targeted by mtr-miR1533a and mtr-miR414c. mtr-miR414h, mtr-miR1533a, mtr-miR1533b, mtr-miR1533c and mtr-miR5021m target WRKY family transcription factor. Squamosa promoter-binding 13A-like protein is targeted by mtr-miR5021a, mtr-miR5021b, mtr-miR5021c and mtr-miR5021d. Homeobox associated leucine zipper protein is targeted by mtr-miR166b, mtr-miR1533b, mtr-miR5021e, mtr-miR5021f and mtr-miR5021m. A network model of the regulation of these transcription factors by miRNAs is depicted in Fig 6, where different miRNAs are shown to target each transcription factors.

**Fig 6:**
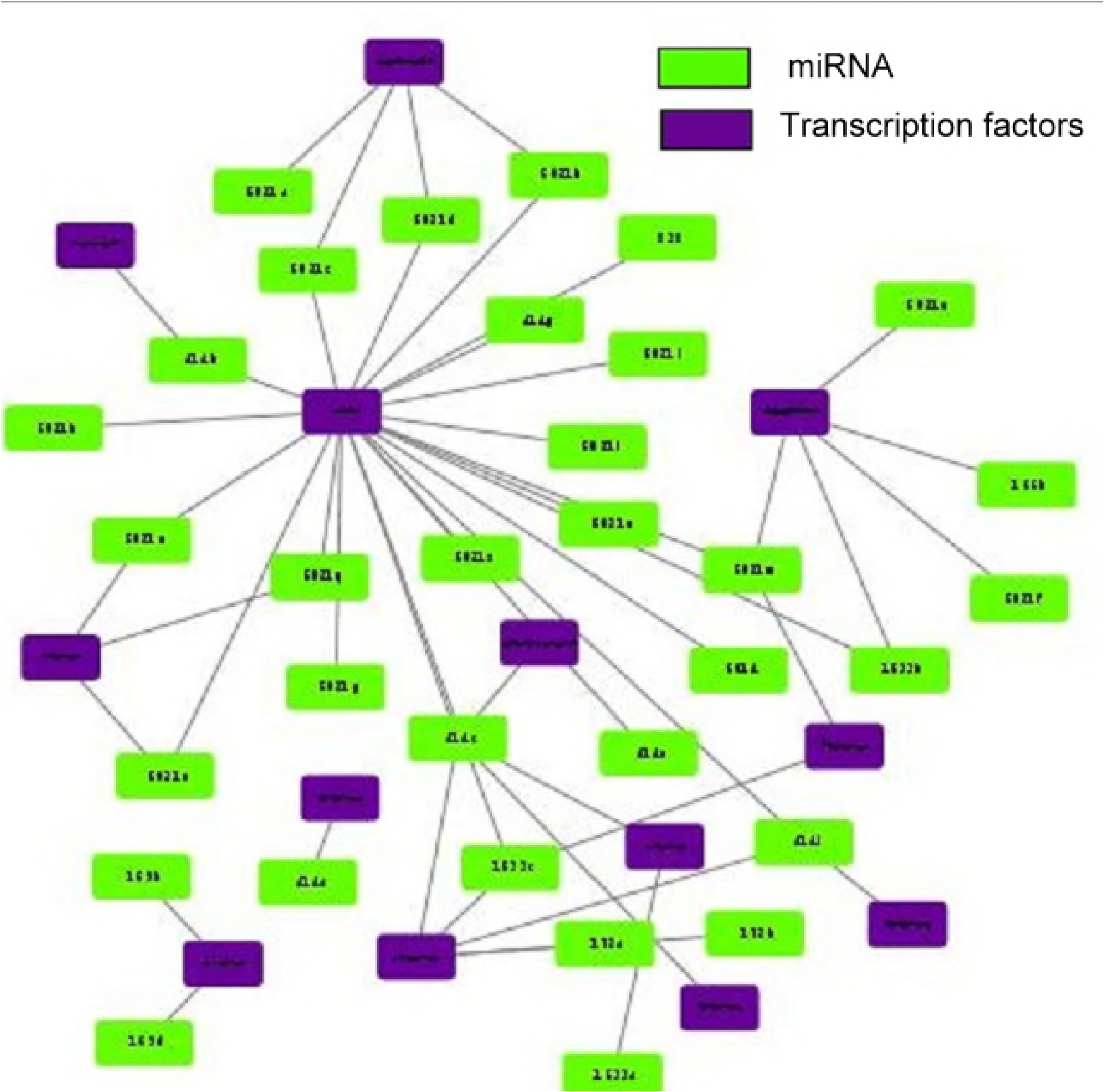
Regulation of transcription factors

Functional annotation of these 621 proteins is carried out using UniProt-GOA (40). The UniProt *M. truncatula* proteome UP000002051 was used to obtain the information about the biological processes, molecular functions and the cellular components where the target proteins were found to be localized. The distribution of the targets involved in these processes is shown in Fig 7 A-C.

**Fig 7:**
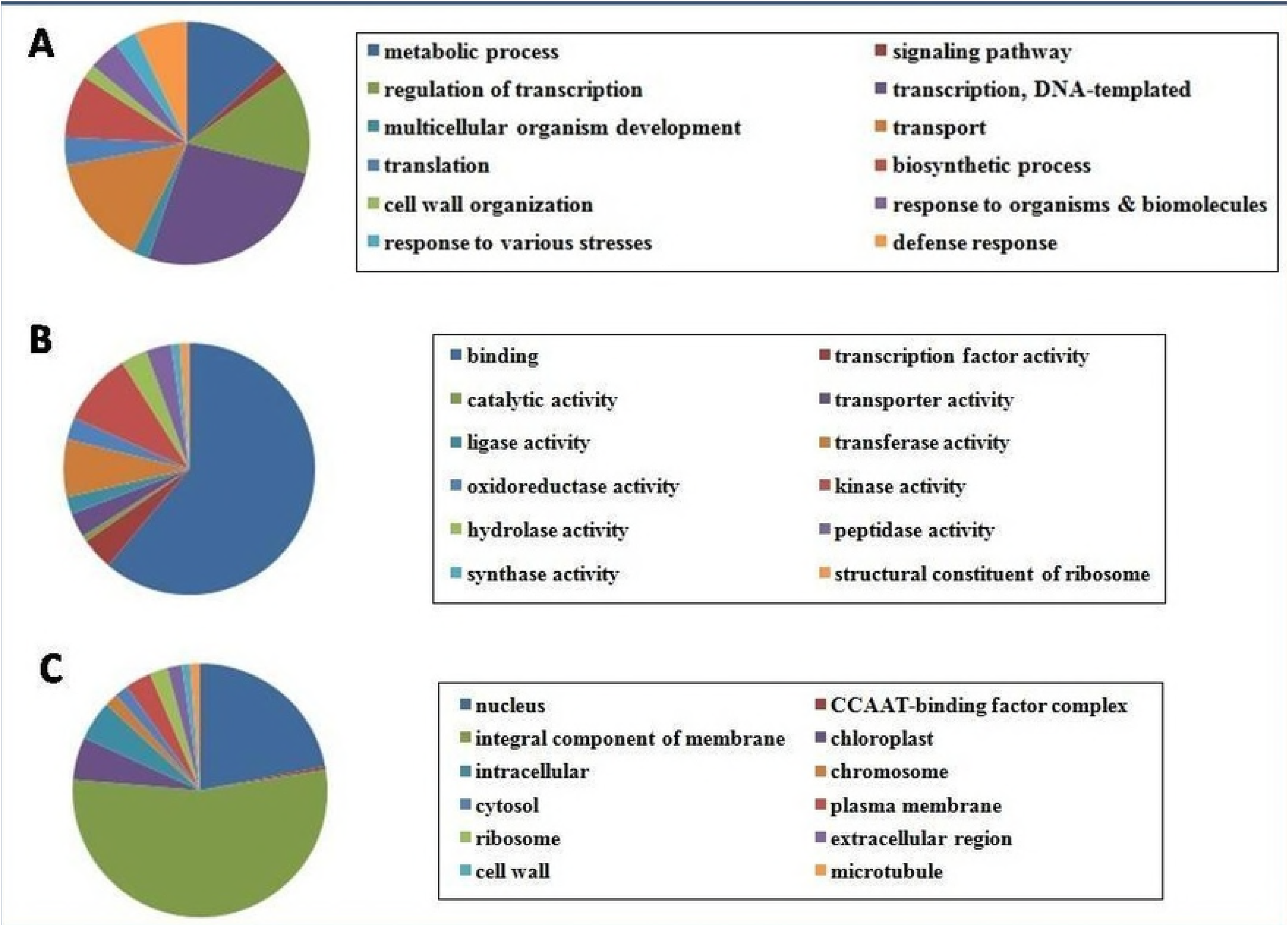
GOannotation of miRNA targets distribution A. biological processes, B. molecular functions, C. cellular components.

Target proteins associated with biological functions shown maximum involvement in transcription (26.3%) and regulation of the transcription processes (13.8%), followed by transport (14.8%) and metabolic processes (13.28%) (Fig 7A). Remaining targets were found to be involved in plethora of biological processes including but not limited to signaling pathways, multicellular organism development, translation, cell wall organization, biosynthetic processes and biotic and abiotic stress responses.

Majority of the targets (61%) associated with molecular functions are found to be involved in binding of various biomolecules including ATP, ADP, DNA, chromatin, along with the binding of different metal ions including Zn and Ca (Fig 7B). Other targets are involved in various enzymatic activities, catalytic activities, transcription factor activities and transporter activities (Fig 7B).

Of the different targets contained in cellular components, majority (54%) are found to be localized in the integral component of the membrane (Fig 7C). Of the remaining, 22% are found in nucleus and others are found to be localized in chromosome, cytosol, plasma membrane, ribosome, cell wall, microtubule and other parts of the cell (Fig 7C).

### Prediction of miRNA targets on lncRNAs

In recent studies lncRNAs have been emerged as an important regulator of transcription and competing endogenous RNAs (ceRNAs) (48). lncRNAs are involved in the regulation of growth, differentiation, stress responses, developmental processes and chromatin modification (49–51). For example, cold induced long antisense intragenic RNA (COOLAIR) and cold assisted intronic noncoding RNA (COLDAIR) are the lncRNAs that are responsible for the suppression of *Flowring Locus C* (FLC) mediated transcription by chromatin modification which in turn regulate reproduction of plants (52,53). lncRNA *P/TMS12-1* is responsible for male fertility in plants (54).

lncRNAs regulate miRNA mediated inhibition by mimicking miRNA targets (55). Hence, it is important to find the miRNA targets on lncRNAs to understand the cross-talks between these two ncRNAs. Previous studies showed that lncRNA Induced by Phosphate Starvation1 (IPS1) is responsible for sequestration of miR399 by forming nonproductive interaction with the complementary miRNA (55).

Among the newly predicted miRNAs, 51 miRNAs target 282 lncRNAs. Details of the miRNA-lncRNA interactions are available in supplementary table S3. Network model built using cytoscape 3.6.0 of this interaction is depicted in Fig 8a. Number of lncRNA targets of miRNAs varies from 1 to 86 with an average of 25 (Fig 8b). About one-third of the miRNAs that target lncRNAs have only one interacting partner. Chromosomal distribution of these lncRNA targets of the novel miRNAs is shown in Fig 8c along with the distribution of the predicted miRNAs.

**Fig 8a:**
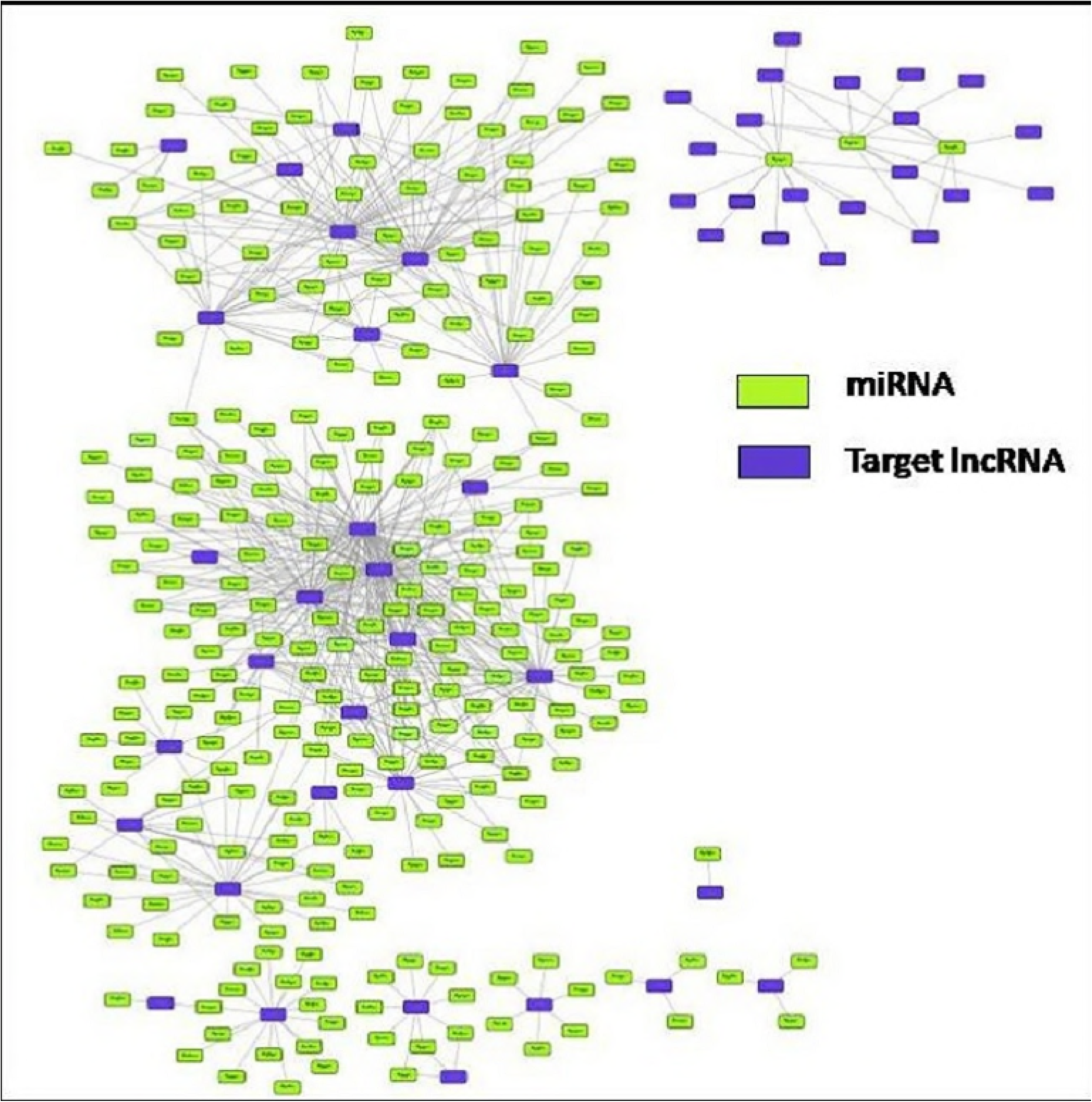
miRNA-lncRNA interaction model

**Fig 8b:**
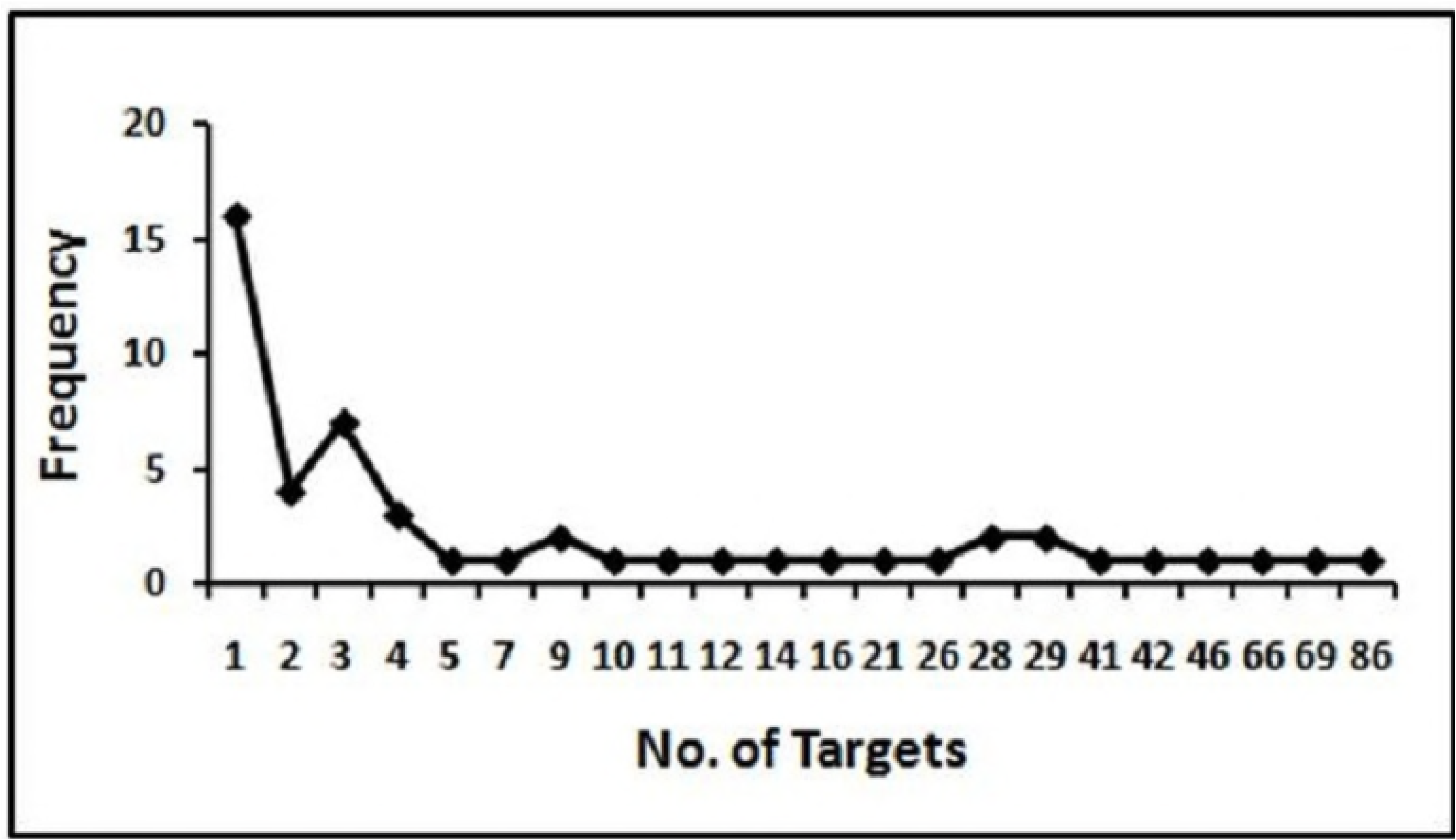
Frequency distribution of miRNA-lncRNA interaction

**Fig 8c:**
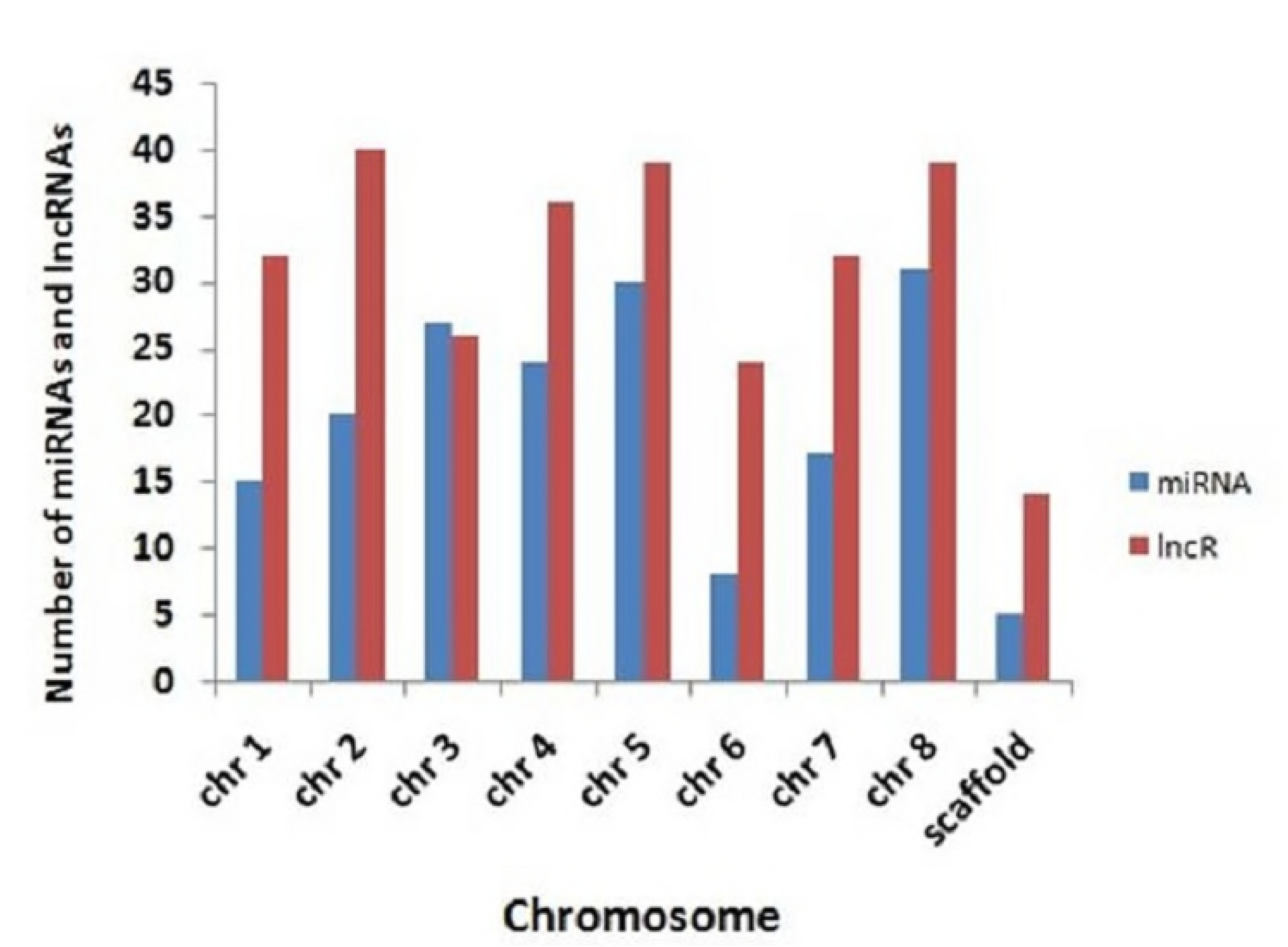
Chromosomal distribution of lncRNA targets

## Conclusion

In this study we have predicted miRNAs of *M. truncatula* and their targets on mRNAs and lncRNAs. A total of 178 mature miRNAs from 40 different miRNA families are predicted; among them 149 are novel and are not reported in miRBase21. We have also predicted 621 characterized target proteins for 93 miRNAs. Among the 149 novel miRNAs, 51 miRNAs target 282 lncRNAs. Many of the novel miRNAs are actively taking part in different stages of symbiotic processes, as well as regulate different transcription factors during post-transcriptional gene regulation (PTGS). *M. truncatula* is an important legume owing to its nitrogen fixation ability and information gained about the miRNAs and their functions related to symbiosis as well as PTGS can provide insight into the gene regulation processes at molecular level in this model plant. Knowledge gained from the role of miRNAs regulating symbiosis in *M. truncatula* will have a positive impact on the nitrogen fixing ability of this model plant which in turn will improve soil fertility.

## LIST OF ABBREVIATIONS

ncRNAs: non-coding RNAs
lncRNAs: long non-coding RNAs
miRNAs: microRNAs
pre-miRNA: precursor miRNAs
MFEI: Minimal Folding Free Energy Index
NQ: normalised Shannon entropy
Npb: normalized base-pairing propensity
ND: normalized base-pair distance
SSR: simple sequence repeat
CDS: coding DNA sequences
PTGS: Post-transcriptional gene silencing

## ETHICS APPROVAL AND CONSENT TO PARTICIPATE

Not applicable.

## AUTHORS’ CONTRIBUTIONS

RPB and JB conceived the study and participated in its design and coordination. MRC and CN performed the study. MRC wrote the manuscript.

## ACKNOWLEDGEMENTS

MRC is thankful to the IIT Kharagpur for research fellowship. Authors thank Nithin C. for providing the computer programs.

